# Cortical Silent Period reflects individual differences in action stopping performance

**DOI:** 10.1101/2020.07.28.219600

**Authors:** Mario Paci, Giulio Di Cosmo, Mauro Gianni Perrucci, Francesca Ferri, Marcello Costantini

## Abstract

Inhibitory control is the ability to suppress inappropriate movements and unwanted actions, allowing to regulate impulses and responses. This ability can be measured via the Stop Signal Task, which provides a temporal index of response inhibition, the stop signal reaction time (SSRT). At the neural level, Transcranial Magnetic Stimulation (TMS) allows to investigate of motor inhibition within the primary motor cortex (M1), such as the Cortical Silent period (CSP). CSP’s length is an index of GABA_B_-mediated intracortical inhibition within M1. Although there is strong evidence that intracortical inhibition varies during action stopping, it is still not clear whether differences in the neurophysiological markers of intracortical inhibition contribute to behavioral differences in actual inhibitory capacities. Hence, we here explored the relationship between intracortical inhibition within M1 and behavioral response inhibition. GABA_B_ergic-mediated inhibition in M1 was determined by the length of CSP, while behavioral inhibition was assessed by the SSRT. We found a significant positive correlation between CSP’s length and SSRT, namely that individuals with greater levels of GABA_B_ergic-mediated inhibition seem to perform overall worse in inhibiting behavioral responses. These results support the assumption that individual differences in intracortical inhibition are mirrored by individual differences in action stopping abilities.

## Introduction

Inhibitory control is a central executive function that allows to temporarily withhold or completely suppress inappropriate or unintended responses, even after these are already initiated. This ability plays a pivotal role in everyday life because behaving in a goal-directed manner constantly requires a quick and efficient regulation of our impulses and responses^19^. Lacking an efficient inhibitory control may result in a number of different dysfunctional behaviors, as evidenced in several medical and psychiatric conditions^39^ such as attention-deficit/hyperactivity disorder^49^, eating disorders^2^ substance abuse disorders^56^ and obsessive-compulsive disorder^47^.

At the behavioral level, one of the most reliable paradigms employed for measuring response inhibition is the Stop Signal Task^37,40,60,63^ (SST). This task allows estimating individuals’ ability to suppress a response already initiated, as it measures the temporal dynamics underlying successful response inhibition^40–42,60,61^. Performance in this task is highly variable across the normal population and reaches abnormally longer values in clinical conditions^39^ including attention-deficit hyperactivity disorder^49^ (ADHD) and obsessive compulsive disorder^47^ (OCD), as well as Gambling Disorder^12^. Hence finding biomarkers of response inhibition is desirable.

At the neural level, Transcranial Magnetic Stimulation (TMS) has been widely employed to investigate the electrophysiological markers of motor inhibition in the brain^19,21,25–28,30,38,50,57^. Different TMS-EMG protocols can be used to measure levels of intracortical inhibition within the primary motor cortex (M1). Specifically, it can be quantified either from the intensity of Short-interval intracortical inhibition (SICI) and Long-interval intracortical inhibition (LICI), obtained with two different paired-pulses procedures or from the length of the cortical silent period (CSP), measured following a particular single pulse procedure^29,51,64^. In particular, the CSP is a cessation in the background voluntary muscle activity induced by a single suprathreshold TMS pulse delivered on M1 during tonic contraction of the target muscle^29,51,54^ (Figure 1). This parameter is obtained by measuring the time interval between the onset of the MEP and the restoration of the muscle activity, and it is expressed in milliseconds (ms). The first part of the CSP (50-75 ms) is thought to be partially due to spinal cord inhibition contributions, while its latter part is entirely mediated by motor cortical postsynaptic inhibition^29,51^. Overall, the length of the CSP is considered an index of the levels of slower inhibitory postsynaptic potentials GABA_B_ inhibition within M1^5,51,80^. Crucially, while SICI and LICI provide an amplitude measure of intracortical inhibition, the CSP provides a temporal measure of this process. Hence, even though both LICI and CSP could be treated as markers of GABA_B_-mediated inhibition, these two measures do not overlap, as they reflect different aspects^34,46^. Specifically, LICI is an electric potential difference expressed in millivolts which represents the magnitude of inhibition, while CSP represents the duration of intracortical inhibition. Given the complementary nature of these different measures, several works have already tried to employ TMS during the SST to probe the fluctuations in the levels of corticospinal and intracortical inhibition within M1 associated with concurrent action preparation and action stopping^20^.

**Figure 1.**
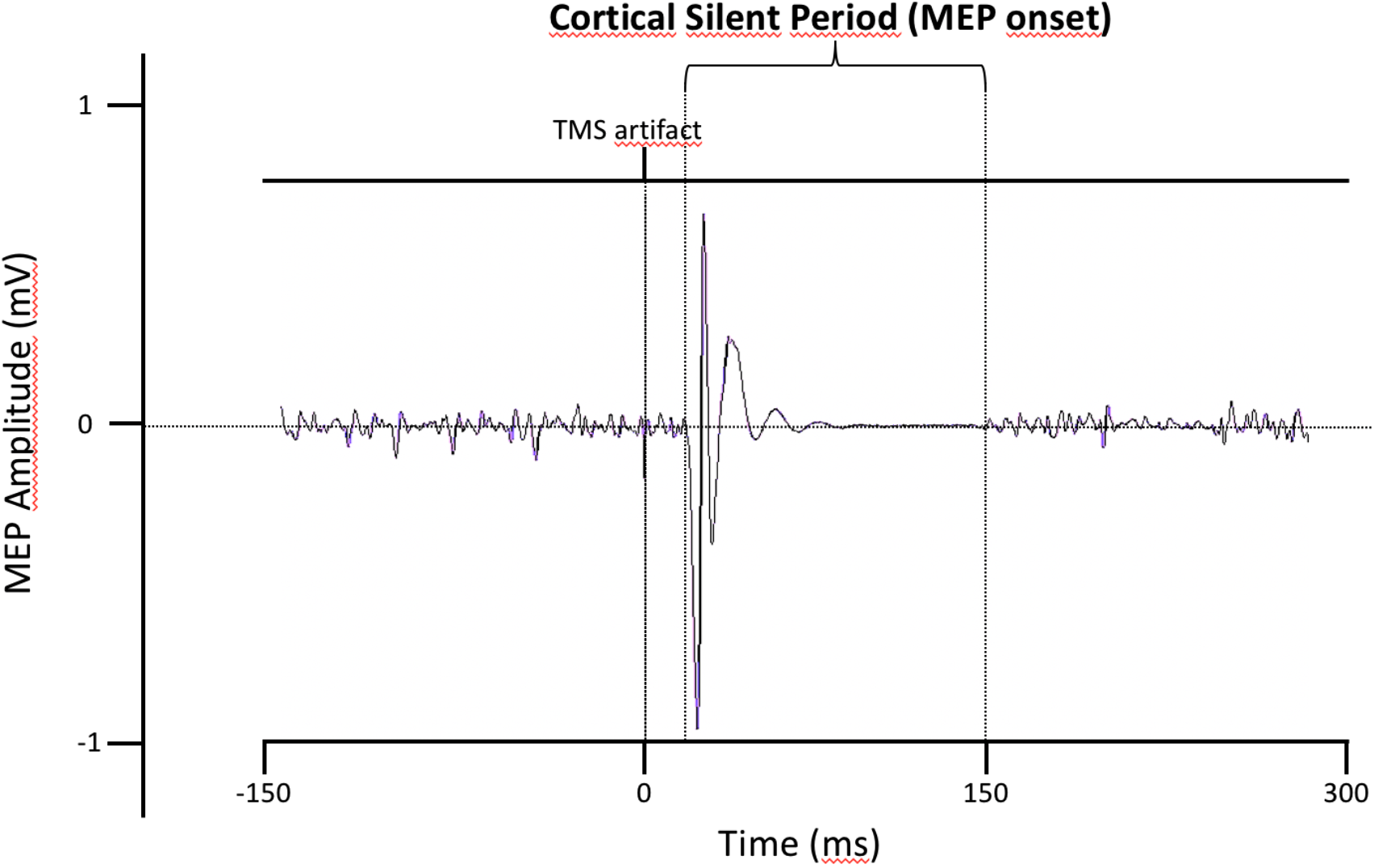
Representative traces of the FDI’s cortical silent period at 30% maximal voluntary contraction (MVC).

Interestingly, studies in which TMS was delivered online on participants during the SST revealed that behavioral motor inhibition is deeply influenced by the ongoing electrophysiological modulation of corticospinal motor excitability and inhibition within M1^20^. For example, in go trials, action preparation induces a significant progressive increase in the levels of corticospinal excitability in the contralateral M1^15,27,43,59^ (≈ 130-175 ms after the onset of the go signal), while during stop trials action inhibition induces both a widespread decrease in corticospinal excitability^15^ (≈ 140ms after the onset of the stop signal) and, concurrently, a significant increase in SICI. Overall, the ongoing modulation of corticospinal excitability and intracortical inhibition within M1 appears to be critical to the successful restraint and cancellation of actions. Nevertheless, it is still unclear whether individual differences in these neurophysiological markers of intracortical inhibition might be related to actual behavioral individual differences in inhibitory control efficiency^12,31^. Recently quite a few studies^7–12,31–33^ have investigated whether and to what extent individual levels of resting-state SICI and LICI measured offline might reflect individual differences in the efficiency of the inhibitory processes, indexed by the length of the Stop Signal Reaction Time (SSRT). Taken together, these studies support the hypothesis that trait-like individual differences in the neurophysiological markers of intracortical inhibition (and SICI in particular) can predict an individual’s actual behavioral motor inhibition capacities^12^. Hence, TMS-derived measures of intracortical inhibition might be effectively employed as biomarkers of motor inhibition performance. However, to date, no studies have investigated whether the CSP’s length measured offline might also candidate as a viable biomarker of motor inhibition, notwithstanding this being the only TMS-based parameter measured as a time interval. This is the aim of the current study.

## Results

### Behavioral data

Raw data were processed via a customized R software (version 3.6.2) for Windows, using the code for the analysis provided by Verbruggen and Colleagues^60^. The average SSRT was 215 ms (SD = 20.6 ms). The table below (Table 1) shows the average and SD of the main measures obtained from the Stop Signal Task: Stop Signal Reaction Time (SSRT), Stop Signal Delay (SSD), RT on go trials (goRT), probability of responding on a stop trial [p(respond|signal)], RT on unsuccessful stop trials (sRT), probability of go omissions (miss), and probability of choice errors on go trials (acc).

**Table 1.** Descriptive statistics of the Stop Signal Task. Values in the table represent means and standard deviations of the descriptive statistics of the Stop Signal Task. Legend: SSRT = Stop Signal Reaction Time; SSD = Stop Signal Delay; goRT = RT on go trials; p(respond|signal) = probability of responding on a stop trial; sRT = RT of go responses on unsuccessful stop trials; miss = probability of go omissions; acc = probability of choice errors on go trials.

| Stop Signal task | Mean ± SD |
| --- | --- |
| SSRT | 215 ms ± 21 |
| SSD | 318 ms ± 117 |
| goRT | 546 ms ± 105 |
| $p(\text{respond}\vert \text{signal})$ | 49% ± 1 |
| sRT | 480 ms ± 89 |
| miss | 2% ± 3 |
| acc | 99.6% ± 0.4 |

### Neurophysiological data

The CSP duration was defined as the time elapsed between the onset of the MEP and the time at which the post-stimulus EMG activity reverted to the pre-stimulus level. The analysis of CSPs was carried out using Signal 6.04 software (Cambridge Electronic Design, Cambridge, UK). Overall, the mean CSP duration was 141 ms (SD = 26 ms). Average relative MEP amplitude was 0.49 mV (SD = 0.29 mV). The average rMT was 55% of the maximum stimulator output (ranging from 48% to 61%, SD = 4%).

### Correlation analysis

The linear correlation between the CSP and the SSRT was significant (Pearson r (27) =.60; *p* =.0009; CI = [0.36 0.77], Figure 2a) and survived robust correlation (skipped Pearson r (27) =.60; CI = 0.36 0.77]). Most importantly, the results of the leave-one-out cross-validation analysis showed a significant correlation between the model-predicted and observed SSRT values (r(27) =.52; *p* =.005, Figure 2b). Moreover, CSP also positively correlated with SSD (Pearson r(27) = −.468; *p* =.014) goRT (Pearson r(27) = −.388; *p* =.046), sRT (Pearson r(27) = −.403; *p* =.037), as well as with the probability of go omissions (miss; Pearson r(27) = −.546; *p* =.003). These correlations were specific for CSP only, as other TMS-derived neurophysiological parameters (rMT, relative MEP amplitude) didn’t correlate with any of the stop signal task significant parameters (Table 2).

**Figure 2a.**
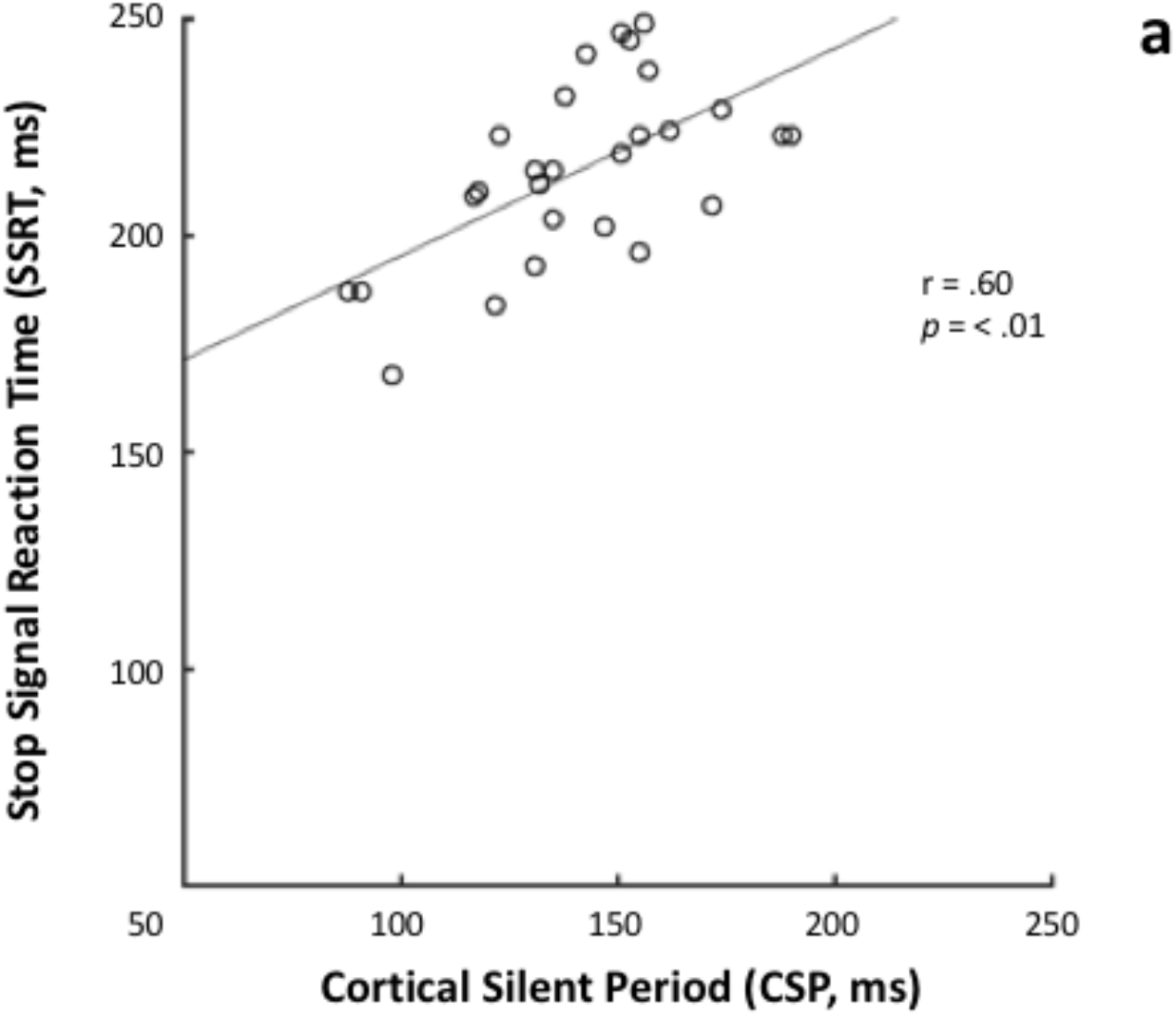
Linear association between cortical silent period (CSP) and Stop Signal Reaction Time (SSRT).

**Figure 2b.**
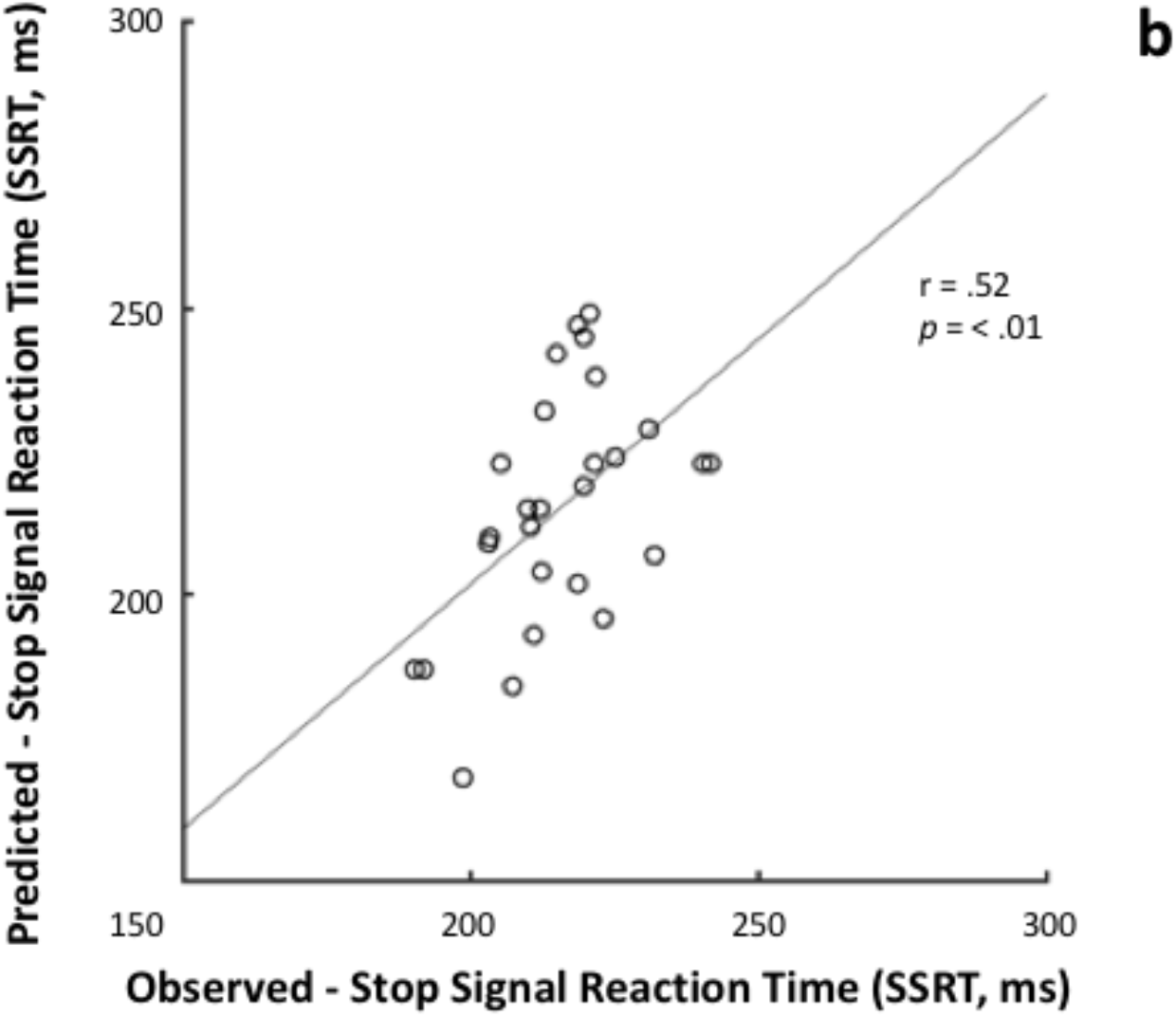
Leave one out cross-validation analysis showing a significant correlation between the model-predicted and observed SSRT values.

**Table 2.**
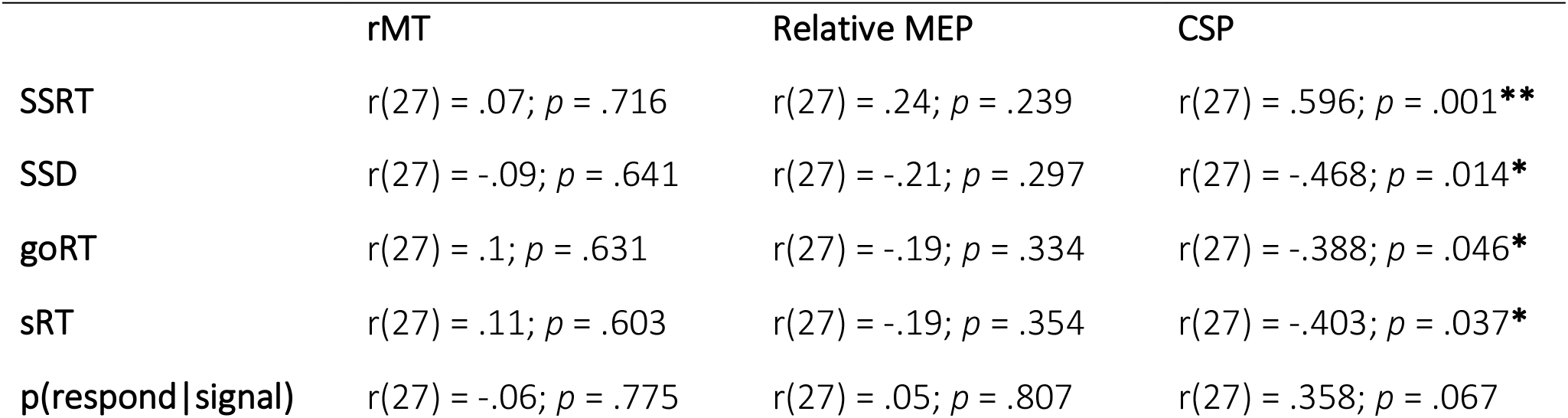
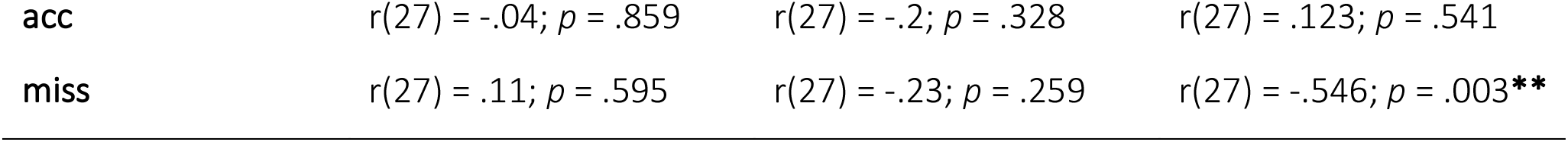
Correlations between the TMS-derived neurophysiological parameters and the stop signal task significant parameters. Values in the table represent Pearson correlational coefficient and significance of the Pearson correlations between each significant Neurophysiological and Behavioral parameter of interest. Legend: rMT = Resting Motor Threshold; Relative MEP = Relative Motor Evoked Potential amplitude; CSP = Cortical Silent Period; SSRT, SSD, goRT, p(respond|signal), sRT, acc = see Table 1; ** = correlation is significant at the 0.01 level (2-tailed); * = correlation is significant at the 0.05 level (2-tailed).

|  | rMT | Relative MEP | CSP |
| --- | --- | --- | --- |
| SSRT | $r(27) = .07$; $p = .716$ | $r(27) = .24$; $p = .239$ | $r(27) = .596$; $p = .001^{**}$ |
| SSD | $r(27) = -.09$; $p = .641$ | $r(27) = -.21$; $p = .297$ | $r(27) = -.468$; $p = .014^*$ |
| goRT | $r(27) = .1$; $p = .631$ | $r(27) = -.19$; $p = .334$ | $r(27) = -.388$; $p = .046^*$ |
| sRT | $r(27) = .11$; $p = .603$ | $r(27) = -.19$; $p = .354$ | $r(27) = -.403$; $p = .037^*$ |
| $p(\text{respond}\vert \text{signal})$ | $r(27) = -.06$; $p = .775$ | $r(27) = .05$; $p = .807$ | $r(27) = .358$; $p = .067$ |
| acc | $r(27) = -.04; p = .859$ | $r(27) = -.2; p = .328$ | $r(27) = .123; p = .541$ |
| miss | $r(27) = .11; p = .595$ | $r(27) = -.23; p = .259$ | $r(27) = -.546; p = .003^{**}$ |

## Discussion

The present study aimed at investigating whether individual differences in the temporal aspect of intracortical inhibition might act as a neurophysiological trait marker reflecting individual response inhibition capacities. Our results revealed a clear relationship between the length of the cortical silent period (CSP) and the stop signal reaction time (SSRT), obtained from the stop signal task (SST). In particular, individuals with longer CSP performed worse at the stop signal task, as indexed by longer SSRT, compared to individuals with shorter CSP. The duration of CSP is a neurophysiological marker of the levels of intracortical inhibition within M1^29,51^. Lengthening of the CSP is observed after disruption of motor attention by sedative drugs such as ethanol or benzodiazepines. Indirect pharmacological evidence supports a largely GABA_B_-mediated origin of the CSP^66–69^. On the other hand, SSRT is a precise index of the duration of the whole chain of processes underlying response inhibition, and so a longer SSRT indicates lower levels of inhibitory control, while a shorter SSRT denotes a better response inhibition. Therefore, our results suggest that CSP might provide a valid trait bio-marker of the quality of action restraint and response inhibition, namely individual inhibitory control capacities.

In general, the relationship between ongoing corticospinal brain activity and behavioral motor functioning has been extensively investigated^20^. Recently, quite a few studies^7–12,31–33^ have investigated the relationship between offline TMS-derived GABA-ergic inhibitory biomarkers (resting-state SICI, LICI) and behavioral motor-inhibitory efficiency. In particular, in their study Chowdhury and Colleagues^12^ showed a negative correlation between individual GABA_A-_ergic intracortical motor inhibition (measured via SICI’s amplitude) and SSRT’s length, indicating that subjects with stronger resting state SICI tend to be faster at inhibiting their responses, and so better at action stopping.

Hence, our results complemented those reported by Chowdhury and Colleagues^12^. Indeed, intracortical inhibition is not modulated through a single process, but it is the synergic result of the interaction between two functionally distinct neural populations mediating either short-lasting ionotropic GABA_A_ postsynaptic inhibition or long-lasting metabotropic GABA_B_ postsynaptic inhibition. Such two mechanisms are indexed by SICI and LICI/CSP respectively^18,45^.

According to the resulting model of interaction between these different cortical inhibitory systems, LICI and CSP do not only inhibit cortical outputs via postsynaptic GABA_B_ receptors, but they even selectively suppress SICI via presynaptic GABA_B_ receptors, causing an overall reduction of GABA release^13,16,55,64^. This GABA_B_ mediated suppression of GABA_A_ effects has been corroborated by converging evidence from in vitro studies^17,18^, as well as pharmacological and TMS studies both at rest and during voluntary movement^46,48,55,64^. Hence, not only SICI and LICI/CSP are mediated by different neural circuits, but the latter can inhibit the first, and so SICI inhibition might result not only from the activity of SICI circuits itself but also as a consequence of increasing GABA_B_-mediated inhibition^55^. Given that CSP is regulated by a GABA_B_-mediated inhibitory network and that longer CSPs are considered an index of upregulated intracortical inhibition^6^, we hypothesize that stronger GABA_B_-ergic circuits, indexed by longer CSP, might unbalance intracortical inhibition during action stopping, resulting in a global reduction of SICI-mediated reactive inhibition and so in a longer SSRT. Interestingly, a similar detrimental effect of CSP’s upregulation has been recently associated with eating disorders, and specifically with Binge Eating Disorder^1^. Keeping this in mind, the negative correlation between SICI’s strength and SSRT found by Chowdhury and Colleagues^12^ and the positive correlation between CSP and SSRT reported here might originate from the same overall inhibitory mechanism. Indeed, it is entirely plausible that these two correlations reflect the opposite effects of these two distinct neural populations mediating different inhibitory sub-process during action stopping, and so that stronger levels of GABA_B_ inhibition might unbalance intracortical inhibition diminishing the GABA_A_-mediated inhibition during action stopping, resulting in worse performance and consequently in a longer SSRT.

Overall, our results suggest that the length CSP measured off-task should be considered as a neurophysiological inhibitory biomarker reflecting individual response inhibition capacities, and specifically that individuals with longer CSP performed worse at action stopping (longer SSRT), compared to individuals with shorter CSP (shorter SSRT). Our results also support the idea that TMS-derived biomarkers might provide a reliable methodology to investigate behavioral individual differences in motor inhibition.

## Methods

### Participants

Twenty-seven 27 (11 males, mean age = 27.84; SD = 3.8; range = 23-38) right-handed naïve participants with normal or corrected-to-normal vision took part in the present study. The sample size of 27 was calculated by using G Power software^23,24^ (v3.1.9.6) analysis assuming an effect size (r) of.62 (based on Chowdhury and Colleagues^12^), an acceptable minimum level of significance (α) of 0.05, and an expected power (1-β) of 0.80.

During the recruitment stage, participants were preliminarily screened for history of neurological disorders and current mental health problems, as well as for hearing and visual difficulties, and completed a questionnaire to check whether they were eligible for a TMS-based study. None of the participants here included reported neither having TMS contraindicators nor having been diagnosed with any psychiatric or neurological disorder, as self-reported. Participants provided their informed consent before taking part in the study. None of the participants reported any negative side effects during or after the TMS procedure. The whole study took place at ITAB (Institute for Advanced Biomedical Technologies) in Chieti and lasted 1 hour and 15 minutes on average. The study was approved by the Ethics Committee of the “G. d’Annunzio” University of Chieti-Pescara and was conducted in accordance with the ethical standards of the 1964 Declaration of Helsinki.

### Measures

#### EMG recording preparation

The surface EMG signal was recorded from the right First Dorsal Interosseous (FDI) hand muscle using three self-adhesive EMG electrodes connected to a CED Micro 1401 (Cambridge Electronic Design, Cambridge, UK). Before electrode placement, recording areas were wiped using an alcohol swab and a pad with abrasive skin prep. Three electrically conductive adhesive hydrogels surface electrodes were then placed along with the target areas of the right hand. Specifically, the positive electrode was placed over the FDI muscle, the negative electrode was placed on the outer side of the thumb knuckle, and the ground electrode was placed on the ulnar styloid process. EMG raw signals were amplified (by a factor of 1000), digitized at a sampling rate of 8 kHz, and filtered using an analogical online band-pass (20 Hz to 250 Hz) and a 50 Hz notch filter. EMG recordings were then stored on a computer for offline analysis with Signal 6.04 software (Cambridge Electronic Design, Cambridge, UK).

#### Transcranial magnetic stimulation

Before TMS administration, participants wore a hypoallergenic cotton helmet which was used to mark the exact location of the FDI hotspot over the left primary motor cortex (M1). Single-pulse TMS was delivered over the left primary motor cortex using a double 70 mm Alpha coil connected to BiStim^2^ Magstim stimulators (Magstim, Whitland, UK) to induce a poster-anterior current flow in the brain. The coil was positioned tangentially to the scalp following the orthodox method^54^, with the handle pointed backward and angled 45 degrees from the midline, perpendicular to the central sulcus. The FDI optimal scalp position for stimulation was identified by maneuvering the coil around the left M1 hand area in steps of 1 cm until eliciting the maximum amplitude motor-evoked potentials (MEPs) in the contralateral FDI muscle using slightly suprathreshold stimuli. Once identified and marked the hotspot on the helmet, coil position was fastened through mechanical support, and its position was constantly monitored by the experimenter. Participants were asked to avoid any head movement throughout the whole TMS session and were also firmly cushioned using ergonomic pads. Afterward, individuals’ resting motor threshold (rMT) was estimated by consistently adjusting the stimulator to find the lowest percentage of the maximum stimulator output necessary to elicit MEPs with a peak-to-peak amplitude of more than 50 **μ**V during muscle relaxation in 5 out of 10 trials^54^. For each participant, rMT was used to determine the specific intensity of TMS suprathreshold stimulation, which was set at 120% of this individual value. This level of stimulation intensity is considered appropriate for studying CSP^24,58^.

### Intracortical Inhibition

Individuals’ CSP was assessed delivering 20 suprathreshold pulses at 120% of rMT while participants were performing an opposition pinch grip at 30% of their FDI’s maximal voluntary isometric contraction (MVC) and maintaining both a static hand posture and a constant level of muscle activity. Individuals’ MVC was used as the reference contraction to normalize the muscular activity between subjects^4,44^ and it was determined by averaging the mean peak-to-peak amplitude of the EMG signal (*μ*V) recorded across three trials lasting 3 seconds each. Before measuring MVC, the experimenter emphasized to participants the importance of performing at their best and of trying to keep the contraction stable during EMG recording. Once determine participants’ MVC, the level of muscular activation was constantly monitored by the experimenter via online data inspection throughout the whole TMS session. Before the TMS session, each participant took part in a preliminary training session to learn how to constantly perform and maintain the appropriate level of FDI contraction (30% MVC) while receiving constant EMG visual feedback displayed on the computer monitor. The TMS session (hotspot mapping procedure, resting motor threshold estimation, and actual experimental session) started only after participants became able to reproduce the adequate level of EMG activity requested without the support of the EMG visual feedback. Each single-pulse TMS stimulation was delivered with an inter-stimulus interval jittered between 8 and 15 seconds to avoid any habituation effect. Trials were rejected if the participant displayed any pronounced head movement before or during the stimulation. For each trial, CSP duration was first quantified as the time between the onset of the MEP and the return of EMG activity to the pre-stimulus level (±2 SD) and then double-checked following a standard procedure^22,58^. CSPs preliminary inspection, analysis, and offline extraction were all carried out using Signal 6.04 software (Cambridge Electronic Design, Cambridge, UK). Remarkably, CSPs were inspected before processing the stop signal task data, and so the inspector was blinded to the relative behavioral results.

### Behavioral level – Motor Inhibition

To measure motor inhibition at the behavioral level we employed the “STOP-IT”^62^, a Matlab and Psychtoolbox^3,35,52^ version of the stop signal task. In the beginning, participants were instructed to place their right hand on a specific site of the computer keyboard (the right index finger over the left arrow key and the right ring finger over the right arrow key) and to maintain this position throughout the whole experiment. The task required participants to perform a speeded response go task while discriminating between two different go stimuli, a left-pointing white arrow, and a right-pointing white arrow, responding to both of them as quickly as possible pressing the left arrow key of the computer keyboard with the right index finger and the right arrow key with the right ring finger, respectively (go trials). However, on 25% of the trials (stop trials), the white go arrow would turn blue after a variable delay (stop signal delay (SSD)), indicating to the participants to withhold their response, whether possible (Figure 3). Crucially, the instructions provided with the task explicitly emphasized that on half of the stop trials the stop signal would appear soon after the go stimulus, making response inhibition easier, while on the other half of these trials the stop signal would be displayed late making response inhibition difficult or even impossible for the participant. Thus, participants were instructed that the task was difficult and that failing in half of the trial was an inherent characteristic of the task itself. They were also instructed to always trying to respond to the go signal as fast and accurately as possible, without withholding their response to wait for a possible stop signal occurrence.

**Figure 3.**
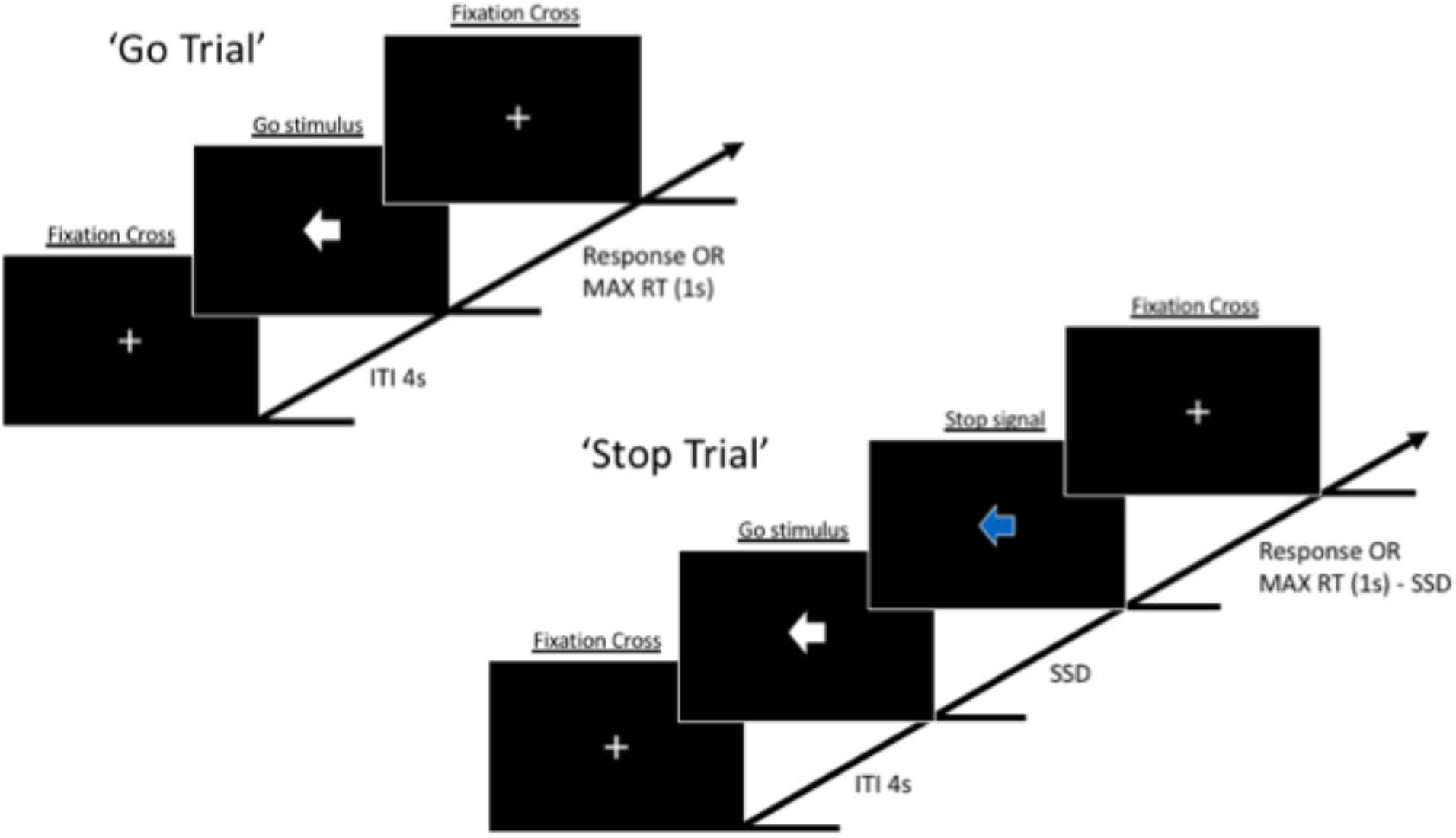
Visual representation of the sequence of events in the stop signal task employed in the present study. On 25% of the trials (stop trials), the white arrow (go signal).

In the go trials, go stimuli lasted 1 second (maximum RT), while in stop trials the combination of the go, the SSD, and the stop signal lasted 1 second in total, with SSD being initially set at 250 ms.

Intertrial interval lasted 4 seconds. Crucially, after each trial the SSD automatically varied in steps of 50 ms as a function of participants previous stop performance, decreasing after each unsuccessful stop trial and increasing after each successful stop, ensuring an overall successful inhibition rate near 50%, and so a p(respond|signal) ≅ .50. The task included an initial practice block providing feedback after each trial. Relevant descriptive statistics obtained from the task included: the probability of go omissions, probability of choice errors on go trials, RT on go trials, probability of responding on a stop trial, Stop Signal Delay, Stop Signal Reaction Time, RT of go responses on unsuccessful stop trials. The Stop Signal data collected from each participant were screened according to the following outlier rejection criteria^14^: (1) probability of responding on stop trials <40% or >60%; (2) probability of go omissions > 25%; (3) probability of choice errors on go trial > 10%; (4) violation of the independent race model (RT of go responses on unsuccessful stop trials > RT on go trials); (5) either negative SSRT or SSRT < 50 ms. These exclusion criteria were adopted because they help to ensure that participants followed the instruction and got engaged throughout the task^14^. In the present study, we used a non-parametric approach for SSRT’s estimation using the integration method with the replacement of go omissions with the maximum goRT^60^. Overall, the whole task comprised 1 practice block of 32 trials (8 stop trials) and 5 experimental blocks of 96 trials (24 stop trials) each. Between blocks, participants were reminded about the instructions and provided with block-based feedback on their performance.

### Procedure

Participants took part in either the TMS session or the behavioral task, administered in random order on the same day with a 10 minutes break between them. The TMS session took place in the TMS/EMG laboratory of ITAB for about 35 minutes, following the same procedure already described above for each participant. The behavioral task took place in one of the Data Collecting Booths of the TEAMLab of ITAB for about 40 minutes. During the behavioral task, participants sat on a comfortable chair in front of a computer monitor with a resolution of 1024 horizontal pixels by 768 vertical pixels, at a distance of approximately 56-57 cm. The tasks were administered on Windows 7 using MATLAB R2016b. the computer monitor refresh rate was 60 Hz. Once finished the two parts, participants were debriefed.

### Data Analysis

To investigate the relationship between behavioral and cortical inhibition we look for a possible correlation between SSRT and CSP.

To control for the specificity of the effect, we run a regression analysis between each of individual stop signal task significant parameters (SSRT, SSD, goRT, sRT, p(respond|signal), acc. ns, miss. ns) and each significant TMS-derived neurophysiological parameters (rMT, amplitude of the accompanying MEP, and duration of CSP). Remarkably, CSPs inspection and extraction were performed before processing the Stop signal task data, and so the inspector was blinded to behavioral results. Moreover, to test the robustness of the relationship we computed skipped parametric (Pearson) correlations^65^ using the Robust Correlation toolbox^53^ and conducted null hypothesis statistical significance testing using the nonparametric percentile bootstrap test (2000 resamples; 95% confidence interval, corresponding to an alpha level of 0.05), which is more robust against heteroscedasticity compared with the traditional t-test^53^. Then, we employed a leave-one-out cross-validation analysis^36^ (i.e., internal validation) to test whether participants’ CSP could reliably predict the SSRT. Specifically, at each round of cross-validation, a linear regression model was trained on n-1 subjects’ values and tested on the left-out participant. Pearson correlations between observed and predicted SSRT values were used to assess predictive power. All statistical tests were two-tailed. To account for the non-independence of the leave-one-out folds, we conducted a permutation test by randomly shuffling the SSRT scores 5000 times and rerunning the prediction pipeline, to create a null distribution of r values. The *p* values of the empirical correlation values, based on their corresponding null distribution, were then computed.

## Data availability

The data analyzed during this study are available from the corresponding author upon reasonable request.

## Author Contributions

M.P., F.F., and M.C. developed the study concept and design. M.P. and M.G.P. developed the experimental setup. M.P. and G.D.C. performed the behavioral data collection. M.P. performed the TMS data acquisition. M.G.P. provided technical support during data acquisition. M.P. analyzed the data. M.P. and M.C. interpreted the data. M.P. wrote the manuscript. All authors contributed to and approved the final manuscript.

## Additional Information

### Competing Interests

The authors declare no competing interests.

## Notes

### Competing Interest Statement

The authors have declared no competing interest.

